# Statistical Methods for Binary Outcomes Adjusting for Outcome Dependent Sampling in Longitudinal Studies with Nonignorable Dropout

**DOI:** 10.1101/2025.08.30.673241

**Authors:** Carter J Sevick, Samantha MaWhinney, Peter L Anderson, Camille M Moore

## Abstract

Longitudinal clinical trials and cohort studies often collect clinical data paired with stored biospecimens. An increasing focus of biomedical research is aimed at leveraging these existing specimens to address new research questions. When a hypothesis of interest proposes to utilize costly, limited or difficult to obtain samples, it may not be possible or desirable to assay all samples. In these situations informed sampling strategies (ISS) can be used to minimize costs and preserve biospecimens by providing a framework to select a subset of subjects that is more informative than a simple random sample. The samples from selected subjects can be assayed and the resulting data can be analyzed in concert with an analytical correction. Dropout is common in longitudinal studies but existing ISS methods do not address nonignorable dropout. Ignoring cases where poor outcomes may influence the propensity to dropout could bias study results. We propose an expansion of current ISS frameworks to account for nonignorable dropout. Mixture models, commonly used to adjust for dropout, are modified to accommodate analysis of data from ISS designs. Methods are available in the BUILD R package.

## 2. Introduction

Longitudinal clinical trials and cohort studies collect valuable clinical data that are often paired with stored biospecimens. An increasing focus of clinical, epidemiological and statistical research is aimed at leveraging these existing data and specimens to address new and important research questions. When the hypothesis of interest proposes to investigate a costly biomarker or utilizes limited and difficult to obtain samples (e.g. cerebral spinal fluid, peripheral blood mononuclear cells), available data can identify a subset of participants whose specimens will be most informative (Schildcrout, 2018a,b; Schildcrout et al., 2015, 2013, 2012; Schildcrout and Heagerty, 2011; Schildcrout and Rathouz, 2010; Schildcrout and Heagerty, 2008; Neuhaus et al., 2014; Neuhaus and Jewell, 1990; Zhou et al., 2007). These sampling schemes are similar to case-control study designs, but instead of using a single outcome, sampling utilizes characteristics of these longitudinal data. Examples of potentially useful characteristics include the sum of binary outcome responses (Neuhaus and Jewell, 1990), outcome summary statistics or distribution (Schildcrout et al., 2013; Zhou et al., 2002), or auxiliary variables (Schildcrout, 2018b; Schildcrout et al., 2012; Zhou et al., 2011).

Informed sampling strategies (ISS), minimize costs and preserve biospecimens by providing methods to select samples of subjects tailored to be more informative about the research question, which can be assayed and analyzed in concert with an analytical correction. However, existing ISS design and analysis methods do not address nonignorable dropout (Schildcrout, 2018a,b). Unfortunately, longitudinal studies, particularly pragmatic trials (Ford and Norrie, 2016) and or those of vulnerable persons, are often plagued by dropout, which if ignored in the sampling design or analysis, may mask associations and bias results (Moore et al., 2017, 2019, 2020; Fairclough, 2010; Daniels and Hogan, 2008; Forster et al., 2011; Little and Rubin, 2014). When the probability of dropout depends on unobserved outcomes even after conditioning on observable data, the data are missing not at random (MNAR) therefore missingness is nonignorable. If MNAR is plausible, non-standard statistical methods and sensitivity analyses should be considered to avoid bias in the results (Little and Rubin, 2014; Daniels and Hogan, 2008; Moore et al., 2017; Forster et al., 2011; Moore et al., 2019, 2020). Outcome dependent sampling (ODS) designs do not intrinsically make any adjustments for MNAR effects and can be biased when applied naïvely (Wilson, 2019). Thus, ISS methods that accommodate dropout are needed. To address these limitations, we propose novel statistical methods for the analysis of ISS in the presence of dropout. We focus on the ODS framework developed by Schildcrout and Heagerty (Schildcrout, 2018a) where the outcome under study is a known longitudinal binary outcome and a categorical exposure must be assessed from stored biosamples. In addition, the nature of the outcome is such that bias from nonignorable dropout is a concern, but is addressable using mixture model methods. To facilitate adoption of ISS methods on a wide scale, we encapsulated our methods for **B**inary outcomes **U**sing **I**nformed sampling strategies for **L**ongitudinal studies with non-ignorable **D**ropout (BUILD) into an R package (BUILD: https://github.com/csevick/build) containing the tools necessary to carry out the analyses in this paper. These tools will save investigators time and resources, while increasing the robustness of statistical inference.

### 2.1 Motivation

Data from the Preexposure Prophylaxis Initiative (iPrEX) Open Label Extension (OLE) study are used as a motivating example. Tenofovir disoproxil fumarate (TDF) or tenofovir alafenamide (TAF) are antiretrovirals used to treat individuals with HIV and also as a Preexposure prophylaxis (PrEP) to prevent new infections in those at risk of acquiring HIV. Concentrations of intracellular tenofovir-diphosphate (TFV-DP) and emtricitabine (FTC-TP) measured in dried blood spots (DBS) have been used to quantify long and short-term adherence, respectively to TDF and TAF regimens (Anderson et al., 2018).

Given multiple spots are collected, typically 5, additional hypotheses may be addressed by assaying residual DBS samples to quantify additional biomarkers of interest. A wide array of DBS assays have been developed, providing straightforward, economical, and non-invasive methods, to quantify a broad range of clinical measures, including drug exposure and/or concentrations (both therapeutic and recreational), environmental exposure biomarkers, multi-omic outcomes (genomic, epigenomic, proteomic and metabolomic), and infectious disease seroprevalence. DBS samples have a finite number of times they may be tested, highlighting the importance of maximizing the efficiency of their use to prolong the viability of the data source for future work. ISS can be instrumental in reducing the number of samples utilized to address future hypotheses in the iPrEX OLE data and more generally.

A secondary research question in the iPrEX OLE study was whether the longitudinal probability of non-compliance, defined as FTC-TP (a short-term adherence biomarker) levels below the limit of quantification (BLQ), was associated with detection of TFV-DP in DBS after the point at which a compliant participant should have reached saturation. While both FTC-TP and TFV-DP data are available for the full iPrEX OLE dataset, we consider a scenario in which FTC-TP levels have already been determined at every measurement point for all participants, while TFV-DP would be obtained by evaluation of stored DBS, creating an opportunity to apply our methodology. Adherence to PrEP has been inversely associated with negative perceptions of side effects and possible stigma resulting from its use (Glidden et al., 2016). Given that these same issues can affect continued study participation, nonignorable dropout is plausible and should be accounted for in analyses.

## 3. Methods

In this section we first review mixture models to account for non-ignorable dropout and methods for outcome dependent sampling. We then introduce our proposed methods which allow the seamless use of ODS while, simultaneously accounting for non-ignorable dropout.

### 3.1 Mixture Models

Mixture model methods account for dropout by factoring the joint outcome-dropout distribution into the dropout time distribution, *h*(*u*), and the distribution of the outcome given dropout, *h*(*y*|*u*). The resulting complete data distribution, *h*(*y*), is / *h*(*y*|*u*)*dH*(*u*). We assume *N* subjects, with known binary outcome *y*_*ij*_ for the *i*^*th*^ subject at time *t*_*ij*_ (*j* = 1, …, *n*_*i*_) with a categorical exposure indexed by *g* = 1, …, *G*. For simplicity, we consider a binary exposure (*g* ∈ {0, 1 }) which can be represented by an indicator variable *X*_*ei*_ = 0, 1 for unexposed and exposed participants, respectively. We consider previous work (Moore et al., 2020) for non-normal outcomes using a generalized linear mixed model (GLMM), with link function *f* (). For binary study outcomes, we assume a logit link function. The conditional model for the outcome given exposure *g*_*i*_ and dropout time *u*_*i*_ is:

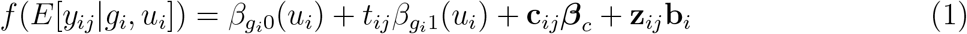

where **c**_*ij*_ is the design vector for covariate effects, ***β***_*c*_, which do not depend on dropout time. **z**_*ij*_ is the design vector for the random effects, *b*_*i*_, where **b**_*i*_ *∼ N* (0, **D**). The *β*_*gip*_ (*u*_*i*_), *p ∈* 0, 1, are exposure group dependent intercept and slope effects, which are functions of dropout time, the parameterization of which could be discrete (a pattern mixture model (Little, 1993)), continuous polynomial (a conditional linear model (Wu and Bailey, 1989)), natural cubic B-splines (Forster et al., 2011; Moore et al., 2019; Moore et al., 2017, 2019, 2020), or penalized splines (Hogan et al., 04a,b). For example, using a conditional linear model with a binary exposure (*x*_*ei*_), would result in the following expression:

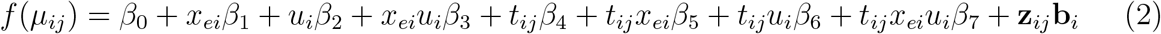

and the dropout time varying intercept and slope would equal *β*_*gi*0_ (*u*_*i*_) = *β*_0_ + *x*_*ei*_*β*_1_ + *u*_*i*_*β*_2_ + *x*_*ei*_*u*_*i*_*β*_3_ and *β*_*g i*1_ (*u*_*i*_) = *β*_4_ + *x*_*ei*_*β*_5_ + *u*_*i*_*β*_6_ + *x*_*ei*_*u*_*i*_*β*_7_, respectively.

When time to dropout is discrete and dependent only on exposure group, marginal parameter estimates may be approximated by a weighed average over the empirical distribution of dropout time (Moore et al., 2020). Let 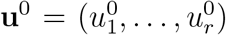 be the *r* unique ordered dropout times of subjects in the dataset, 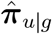 the associated vector of proportions of participants associated with each dropout time, in exposure group *g*, and 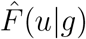 the empirical distribution of dropout time in group *g*. The *p*^*th*^ estimated marginal effect for the *g*^*th*^ exposure group will be:

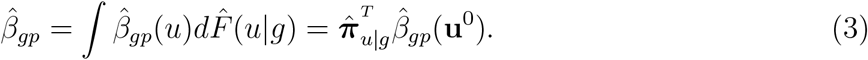

The estimated covariance matrix may be used to construct variances for the marginal parameters, although an adjustment for the estimation of dropout time proportions is required. The Delta Method may be used for this purpose (Hogan et al., 04a,b; Hedeker and Gibbons, 2006; Fairclough, 2010; Sevick et al., 2023), however resampling methods (Davison and Hinkley, 1997; Shao and Tu, 1995) may provide a more robust solution.

### 3.2 Outcome Dependent Sampling (ODS)

#### 3.2.1 Sampling Strategy

In this section, the case-control extension proposed by Schildcrout and Heagerty (Schildcrout, 2018a) is described. A data repository with *N* participants is assumed with longitudinal binary outcomes 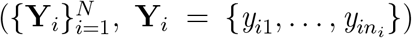 and observed covariate data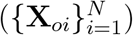. The exposure indicator, to be determined in the sample, is 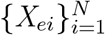. The design matrix of the fixed effects for participant *i* is **X**_*i*_ and **X**_*ij*_ represents the covariate vector for participant *i* at time *j*, which would include *X*_*ei*_, *t*_*ij*_, **c**_*ij*_ and any relevant time by covariate or time by exposure interactions. The vector ***θ*** is adopted to represent parameters associated with the design matrix.

Subjects are first grouped into strata, based on the sum of their longitudinal outcomes. The strata, *V* with *v*_*i*_ being the realized strata assigned to the *i*^*th*^ participant, are defined by:

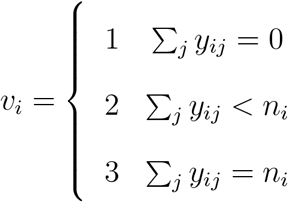

Let *N*^*s*^ be the total number of participants sampled and 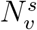 be the number of participants sampled from stratum *v*. A particular sampling design may be specified by the tuple 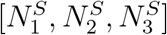.The probability of selection, given outcome stratum assignment is *π*(*v*_*i*_) = *pr*(*S*_*i*_ = 1|*V*_*i*_ = *v*_*i*_), where *S*_*i*_ is a sampling indicator. To maximize efficiency for estimation of time-varying predictors, including time, and their interactions one ISS design approach is to deliberately over sample stratum 2 subjects.

#### 3.2.1 Analysis of ODS designs

Extensions of generalized linear models are used to model binary longitudinal data, including marginalized transition and latent variable models (mTLV) (Schildcrout and Heagerty, 2007; Diggle et al., 2002). Utilizing likelihood based approaches, allows correction for the non-representative sampling using three common approaches: 1) Ascertainment-Corrected Maximum Likelihood (ACML), 2) Inverse probability of selection weighted likelihood (WL) and 3) multiple imputation (MI).

1. ACML: In this method the sampling function is derived conditionally on selection and a marginal expression is arrived at using an application of Bayes’ theorem. The likelihood is,

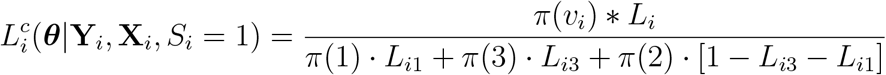

where, *L*_*i*_ = *L*_*i*_(***θ***|**Y**_*i*_, **X**_*i*_) = *pr*(**Y**_*i*_|**X**_*i*_; *θ*) and, where *L*_*i*1_ and *L*_*i*3_ are the likelihood contributions under random sampling had subject *i* been assigned to strata 1 or 3 (all events, or no events) (Mercaldo, 2017).
2. WL: The log-likelihood contributions of each participant are combined in a weighted sum using the inverse of the probability of selection as the weight (Rabe-Hesketh and Skrondal, 2006; Robins et al., 1994; Manski and Lerman, 1977). The weight, *π*(*v*_*i*_)^*−*1^, may be interpreted as the number of participants in the original data source represented by the sampled participant *i*. The likelihood equation, used in conjunction with maximum likelihood (ML) methods for parameter estimation, is:

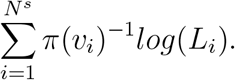 Note that the estimated covariance matrix from the maximization will no longer be valid for inference. To obtain correct inference, robust variance estimates may be used (Rabe-Hesketh and Skrondal, 2006).
3. Multiple imputation (MI): Using a model computed from the sampled participants (now with a known exposure) the exposure is imputed for non-sampled participants, resulting in more efficient use of available cohort data (Schildcrout, 2018a). A additional strength of this method is that standard analytic methods may be used and results synthesized for interpretation with well established procedures (Van Buuren, 2018).

### 3.3 BUILD: Mixture Models to Account for Dropout in ODS Designs

We now expand on the case-control extension proposed by Schildcrout and Heagerty (Schildcrout, 2018a) to accommodate informative dropout using a mixture model approach. As before, subjects are first grouped into strata based on the sum of their longitudinal outcomes, however we propose to augment this stratification with dropout time to ensure representative capture of participants across the range of the dropout distribution. The outcome and dropout strata of the *i*^*th*^ participant, *V*_*i*_ and *U*_*i*_ respectively, are defined as in Table 1

**Table 1.** BUILD Sample Framework: Combined Outcome and Dropout Strata.

| Outcome strata | Dropout strata |  |  |  |
| --- | --- | --- | --- | --- |
|  | 1 | 2 | ... | r |
| 1: $\sum_j Y_{ij} = 0$ | $\pi(1, 1)$ | $\pi(1, 2)$ | ... | $\pi(1, r)$ |
| 2: $0 < \sum_j Y_{ij} < n_i$ | $\pi(2, 1)$ | $\pi(2, 2)$ | ... | $\pi(2, r)$ |
| 3: $\sum_j Y_{ij} = n_i$ | $\pi(3, 1)$ | $\pi(3, 2)$ | ... | $\pi(3, r)$ |

where the *π*(*v*_*i*_, *u*_*i*_) = *pr*(*S*_*i*_ = 1|*V*_*i*_ = *v*_*i*_, *U*_*i*_ = *u*_*i*_) are the sampling probabilities for each stratum. The three methods used to adjust for the non-representative sampling are adapted to this expanded framework. The design matrix for the fixed effects, **X**_*i*_, is augmented to include the necessary effects to define the functions of dropout time, *β*_*gip*_(*u*_*i*_), from the mixture model. Specification of the sampling plan remains as *N*^*s*^ as the total sample size, 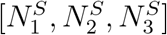 for the marginal outcome strata sample sizes but we further introduce 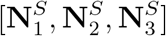 to indicate that outcome-dropout specific sample sizes should be considered.

#### 3.3.1 Ascertainment-Corrected Likelihood

The Ascertainment-Corrected Likelihood (Schildcrout, 2018a) is arrived at by considering the likelihood of the *i*^*th*^ individual conditional on the probability of selection (ascertainment) by application of Bayes’ rule (Casella and Berger, 2002). In this expansion we further condition on dropout strata. The likelihood is:

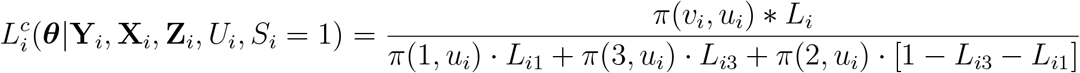

where, *L*_*i*_ = *L*_*i*_(***θ***|**Y**_*i*_, **X**_*i*_**Z**_*i*_, *U*_*i*_) = *pr*(**Y**_*i*_|**X**_*i*_, **Z**_*i*_, *U*_*i*_, ***θ***) and, where **X**_*i*_ is the design matrix for the fixed effects, ***θ*** is the parameter vector for both fixed and random effects. *L*_*i*1_ and *L*_*i*3_ are the likelihood contributions under random sampling had subject *i* been assigned to strata 1 or 3 (all events or no events) (Mercaldo, 2017). In this application, we are confining the ascertainment adjustment to the outcome strata, but leaving the dropout as a conditionally specified aspect of the model. If selection is not proportional across dropout strata then robust variances will need to be computed for valid inference (Rabe-Hesketh and Skrondal, 2006).

#### 3.3.2 Inverse Probability of Selection Weighted Likelihood

The sampling design may also be corrected for by weighting by the inverse probability of selection (Rabe-Hesketh and Skrondal, 2006; Robins et al., 1994; Manski and Lerman, 1977). Let *w*(*v*_*i*_, *u*_*i*_) = *π*(*v*_*i*_, *u*_*i*_)^*−*1^ and *l*_*i*_ = *log*(*L*_*i*_) then, in the inverse weighted likelihood method, the equation for the loglikelihood becomes:

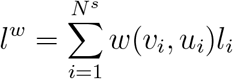

#### 3.3.3 Multiple Imputation

To develop an imputation model we consider the probability of exposure, conditional on outcomes, dropout time and the known covariates, in the population. To the notation developed in section 3.3 we associate parameter vectors ***θ***_*e*_ and ***θ***_*y*_ to the outcome model (for exposure and non-exposure covariates, respectively) and ***θ***_*m*_ to a model for the marginal (with respect to the outcomes) probability of exposure. Utilizing Bayes theorem we have that the probability of exposure, conditional on outcomes, dropout time and known covariates, may be computed in the following way:

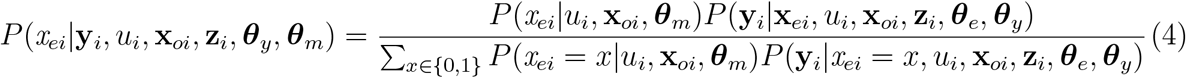

*P* (**y**_*i*_|**x**_*ei*_, *u*_*i*_, **x**_*oi*_, **z**_*i*_, ***θ***_*e*_, ***θ***_*y*_) is the participant’s likelihood contribution.

*P* (*x*_*ei*_|*u*_*i*_, **x**_*oi*_, ***θ***_*m*_) is the marginal probability of exposure, given relevant known covariates and dropout time.

To apply MI in the BUILD setting we recommend a bootstrap strategy (Van Buuren, 2018) in order to incorporate appropriate variability into the parameter estimation. Parameter estimates for equation 4 using BUILD ACML or WL, for the outcomes model, and logistic regression corrected for the sampling design, for the marginal probability of exposure. For the desired number of imputations:

- Draw a stratified bootstrap sample (of size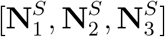, with replacement) from the sampled data (with the assessed exposure)
- Using the bootstrap sample: estimate the parameter vectors for the outcome 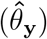 and marginal exposure 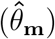 models
- Using the estimates from the previous step, estimate the conditional exposure probabilities in the full data participants (equation 4)
- Impute the exposure in the full data by taking a random draw from a Bernoulli distribution with *P* set to the estimate from the previous step, for all participants
- Analyze the imputed data using a standard mixture model to adjust for dropout effects

Finally, the estimates from each iteration can be combined using Rubin’s rules (Little and Rubin, 2014) to summarize the results.

#### 3.3.4 Marginal Estimates

Marginal parameter estimates are obtained from the adjusted mixture model in the same way as 3.1; however the weights need to be specific to the target population, not the sample. Let *U*_*a*_, *V*_*b*_ and *G*_*c*_ be sets of participants with the *a*^*th*^ ordered dropout time, *b*^*th*^ outcome strata and *c*^*th*^ exposure group, respectively. The joint probability function of these sets is:

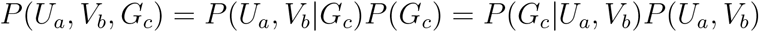

and this implies that 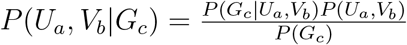.

*P* (*U*_*a*_, *V*_*b*_) can be estimated using the empirical distribution of *U* and *V* using the full study population, while *P* (*G*_*c*_|*U*_*a*_, *V*_*b*_) must be estimated in the sample after exposure status has been assessed. Next, an estimate of *P* (*G*_*c*_) is:

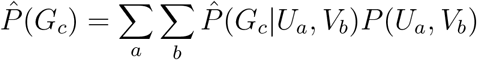

and an estimate of the needed weight is:

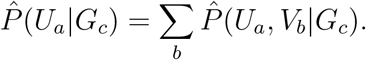

If we use a design where samples of fixed size are drawn from each stratum then a simplification is possible. In this case the sum of participant sampling weights from a given stratum will equal the stratum size in the population. An equivalent computation of the marginalization weight is then:

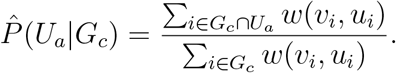

For marginal estimates in BUILD, we set:

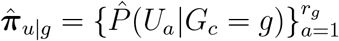

and use equation 3 with the dropout specific parameter estimates from BUILD ACML, WL or MI methods.

## 4. Simulation

### 4.1 Simulation Methods

#### 4.1.1 Data Generation

To study the performance of our methods we extended simulation methods that have been used to study the performance of mixture models for non-ignorable missing data (Hogan et al., 04a; Forster et al., 2011; Moore et al., 2017). Data were simulated from a generalized linear mixed model with a Bernoulli distribution and logit link to represent a data repository consisting of 5000 participants. Longitudinal data consisted of a baseline measure and additional follow up measures, either 5 or 10, with time scaled to be from 0 -

- The simulation model was constructed as a conditional linear model with the form:

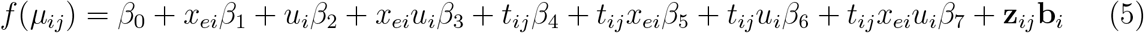

where exposure (*x*_*ei*_) was set to be positive 50% of the time, *u*_*i*_ was the last observed time period for participant *i* and *t*_*ij*_ was the *j*^*th*^ study period for participant *i*. Simulation parameters were set to: *β*_0_ = 1.39 (representing an 80% outcome rate at baseline among the unexposed), *β*_1_ = 0, *β*_2_ = 0, *β*_3_ = 0, *β*_4_ = 0, *β*_5_ = 0.5, *β*_6_ = *−*4, *β*_7_ = 0 or *−*1 and a random intercept variance of 2. Relating this back to equation 1, we have: *β*_*gi*0_ (*u*_*i*_) = *β*_0_ and *β*_*gi*1_ (*u*_*i*_) = *x*_*ei*_*β*_5_ + *u*_*i*_*β*_6_ + *x*_*ei*_*u*_*i*_*β*_7_. Dropout began at the 3rd visit as a beta binomial random variate with *p ∼ Beta*(1, 1). Data for each condition were simulated 500 times.

#### 4.1.2 Analysis

Each simulated data repository was sampled using marginal outcome strata (ODS) and using the expanded outcome by dropout strata (BUILD). All models were estimated as generalized linear mixed models with a logit link, binomial distribution, a single random intercept, and included parameters for time, exposure status, and an interaction for time by exposure, while BUILD models included additional parameters to estimate the dropout effects. Correction for the sampling design was by Ascertainment-Corrected Maximum Likelihood (ACML), weighted likelihood (WL) and multiple imputation (MI). The R optim function (R Core Team, 2023) was used to optimize the likelihood with the BFGS method. Integration of the likelihood function was by Gauss-Hermite quadrature (McCulloch et al., 2008) with 10 points. Analysis by MI was carried out with 30 imputations. Robust variances were computed for WL (Rabe-Hesketh and Skrondal, 2006), while model based were used for ACML and MI. Sampling rates were set to give priority of estimation to change over time (50 never, 700 some, 50 always). The percentages of sampling units within each ODS strata for a typical simulation were [10.4, 59.5, 30.1].

Table 2 presents the percentages of participants in ODS strata by dropout time, typical in a simulation with 5 follow up visits. When applying BUILD methods, the marginal sampling cells were divided proportionally across dropout strata, rounding to the nearest whole number. If the computed proportion to be sampled was between zero and one, exclusive, then one individual was selected.

**Table 2.** Percentage of simulated participants within ODS strata, by dropout time.

| Time point | 0.2 | 0.4 | 0.6 | 0.8 | 1 |
| --- | --- | --- | --- | --- | --- |
| Never | 2.6 | 1.8 | 1.7 | 1.7 | 2.5 |
| Some | 5.8 | 9.0 | 13.0 | 15.2 | 16.5 |
| Always | 11.7 | 8.9 | 5.5 | 3.2 | 0.8 |

#### 4.1.3 Performance Evaluation

A total of 4 conditions were studied (combinations of 5 and 10 follow up visits and a dropout effect on the time by exposure interaction vs none), on 6 different models (ODS ACML, weighted likelihood ODS (WL ODS), BUILD ACML, BUILD weighted likelihood (WL BUILD) and multiple imputation (MI ODS and MI BUILD)). Five hundred simulations were run per condition and we evaluated the performance of the 6 models in terms of bias, coverage of 95% confidence intervals (CI) and mean square error (MSE) for all parameters in the ODS only models compared to marginal parameter estimates from BUILD.

### 4.2 Simulation Results

In general, BUILD models showed less bias and coverage rates closer to nominal than the ODS counterparts (Table 3). Parameter estimates from ODS tended to have less variability, however, MSE estimates were greater, and coverage probability was lower, than BUILD to the extent that parameters were affected by non-ignorable dropout. Specifically, when the dropout effect was large, as in the slope estimates for exposed and control, BUILD was markedly superior in all three metrics. However, when the dropout effect was slight and the parameter was time varying (time:exposure), bias and coverage remained much too extreme for ODS but MSE was actually smaller than BUILD (Table 3). It is interesting to note that when the time:exposure interaction was not directly affected by dropout BUILD remained superior in bias and coverage, even though lagging behind in terms of MSE (see supplementary Table S1, available online). Although, simulated intercept terms did not depend on dropout time, marginal intercept terms were better estimated by BUILD, but intercept shifts (exposure) were better estimated by ODS (Table 3). This could be due to the intercept and time:exposure parameters being indirectly affected by the effect on time while exposure had no direct or indirect effect of dropout. When choosing which effects to correct for dropout it would be advisable to consider indirect effects that may occur through an interaction effect, but to not correct estimates that have no theoretical relationship with dropout.

**Table 3.** Simulation Results: BUILD vs. ODS, Dropout Effect on Time and the Time by Exposure Interaction. Results based on 500 simulations of BUILD designs with marginal outcome strata sizes of [50, 700, 50] with proportional distribution of the sample over dropout time. Dropout was uniform over study time, beginning at visit 3. 30 imputations used for MI.

| Effect | Target value |  | Method | Bias |  | Coverage |  | MSE |  |
| --- | --- | --- | --- | --- | --- | --- | --- | --- | --- |
|  | N Follow Up |  |  | N Follow Up |  | N Follow Up |  | N Follow Up |  |
|  | 5 | 10 |  | 5 | 10 | 5 | 10 | 5 | 10 |
| Intercept | 1.39 | 1.39 | ACML BUILD | 0.003 | 0.012 | 98 | 98.2 | 0.019 | 0.012 |
|  |  |  | ACML ODS | 0.01 | 0.064 | 97.4 | 95 * | 0.016 | 0.016 |
|  |  |  | MI BUILD | 0.002 * | 0.005 * | 92 | 94.4 | 0.015 * | 0.009 * |
|  |  |  | MI ODS | 0.14 | 0.139 | 79.6 | 69.8 | 0.035 | 0.029 |
|  |  |  | WL BUILD | 0.004 | 0.006 | 98.2 | 99 | 0.018 | 0.012 |
|  |  |  | WL ODS | 0.141 | 0.143 | 95.6 * | 90.4 | 0.039 | 0.033 |
| Exposure | 0 | 0 | ACML BUILD | 0.005 | 0.007 * | 94.6 | 95.3 | 0.046 | 0.029 |
|  |  |  | ACML ODS | 0.003 * | 0.02 | 95.2 | 94.6 | 0.036 * | 0.026 * |
|  |  |  | MI BUILD | 0.005 | 0.009 | 92.8 | 94.2 | 0.054 | 0.033 |
|  |  |  | MI ODS | 0.03 | 0.036 | 94.2 | 94.2 | 0.058 | 0.035 |
|  |  |  | WL BUILD | 0.004 | 0.008 | 95 * | 95.4 | 0.055 | 0.034 |
|  |  |  | WL ODS | 0.034 | 0.035 | 94.4 | 95.2 * | 0.059 | 0.036 |
| Slope in control | -2.40 | -2.20 | ACML BUILD | 0.007 | 0.028 | 93.8 | 94.9 * | 0.11 | 0.107 |
|  |  |  | ACML ODS | 0.941 | 1.099 | 1.4 | 0 | 0.934 | 1.244 |
|  |  |  | MI BUILD | 0.002 * | 0.009 * | 92.2 | 94.4 | 0.076 * | 0.064 * |
|  |  |  | MI ODS | 1.056 | 1.145 | 0 | 0 | 1.151 | 1.334 |
|  |  |  | WL BUILD | 0.008 | 0.024 | 95.8 * | 96 | 0.117 | 0.109 |
|  |  |  | WL ODS | 1.056 | 1.149 | 0.8 | 0 | 1.177 | 1.362 |
| Slope in exposed | -2.50 | -2.25 | ACML BUILD | 0.019 | 0.004 * | 95.8 * | 95.5 * | 0.103 * | 0.109 * |
|  |  |  | ACML ODS | 1.135 | 1.357 | 0 | 0 | 1.345 | 1.88 |
|  |  |  | MI BUILD | 0.018 * | 0.01 | 96.8 | 96.4 | 0.113 | 0.111 |
|  |  |  | MI ODS | 1.245 | 1.409 | 0 | 0 | 1.592 | 2.009 |
|  |  |  | WL BUILD | 0.018 * | 0.01 | 96.8 | 96.4 | 0.113 | 0.111 |
|  |  |  | WL ODS | 1.258 | 1.419 | 0 | 0 | 1.652 | 2.055 |
| time:exposure | -0.10 | -0.05 | ACML BUILD | 0.015 * | 0.035 * | 93.8 | 94.9 * | 0.222 | 0.212 |
|  |  |  | ACML ODS | 0.194 | 0.258 | 89.8 | 86.6 | 0.14 * | 0.141 * |
|  |  |  | MI BUILD | 0.016 | 0.042 | 93.4 | 93.8 | 0.258 | 0.231 |
|  |  |  | MI ODS | 0.189 | 0.264 | 90.8 | 87 | 0.163 | 0.147 |
|  |  |  | WL BUILD | 0.016 | 0.041 | 94.4 * | 95.8 | 0.272 | 0.239 |
|  |  |  | WL ODS | 0.202 | 0.27 | 89.6 | 87.2 | 0.185 | 0.154 |

*Note:*
\* Best performing metric for the group.

When considering performance among the 3 methods within BUILD only, the performance of the three methods was mixed and there was no method that was uniformly better (Table 3). Differences between the estimates were never prohibitively large, and this, coupled with the greater complexity of both ACML and MI and the need to specify an additional model for exposure with MI (which could be an additional source of error) leads to our favoring of weighted likelihood for most applied circumstances. In cases where the full population data are available and there are additional variables that do not make sense for the outcome model, but may lead to a more informative imputation model, MI could be given more weight in the decision process.

## 5. Application

### 5.1 Data

Data for this example were drawn from the iPrEX OLE (Grant et al., 2014) study on adherence to pre-exposure prophylaxis (PrEP) in an HIV negative (at baseline) cohort of men who have sex with men (MSM) and transgender women at risk of HIV infection. In the full data, 1603 participants were identified from prior randomized placebo controlled trials on efficacy of PrEP and were enrolled between June 2011 and June 2012. Participants were offered daily oral FTC/TDF PrEP at enrollment. DBS were scheduled to be collected at 4, 8, 12 (and then every 12 up to 72 weeks) weeks after initiation of PrEP.

### 5.2 Methods

For this analysis we focused on participants with at least 2 measurements from week 11 to 77, resulting in 953 participants with 3242 measurements. Baseline was considered to be the first record in this time frame. Dropout was assumed after the last observed measurement time, and was grouped as 17-28, 29-41, 42-53, 54-65 and 66+ weeks. Based on similarity of FTC-TP BLQ (the outcome of interest) trajectories, two dropout strata were formed: last measure from 17-41 weeks vs. 42+ weeks (Figure 1). Outcome strata (never, sometimes, always) contained 368, 365 and 220 participants with 9.5%, 12.6% and 18.6% early dropout, respectively.

**Figure 1.**
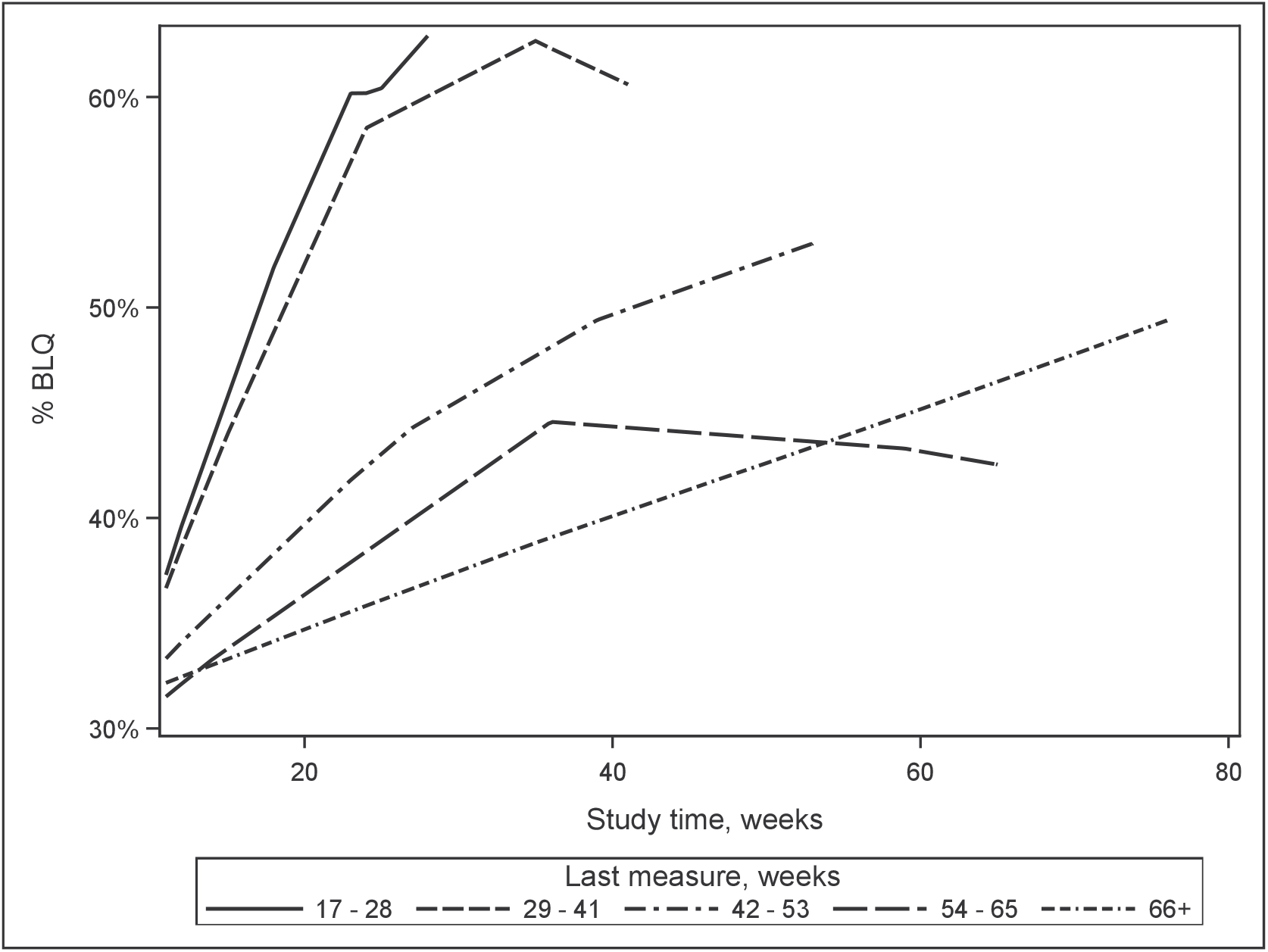
Observed trajectories in undetectable FTC-TP across study measurement periods, by last measure time, for all participants

To adjust for informative dropout, a pattern mixture model was constructed with a single indicator for early dropout vs. late dropout/complete ascertainment. Other covariates included time from steady state baseline (scaled to show the estimated effect over 1 year), whether the first DBS TFV-DP was BLQ, as well as an interaction between the two. An interaction term for each covariate with the early dropout indicator was added. A random intercept model was chosen to account for multiple measurements per patient.

Since interest centered on comparing trends over time, we decided to over sample outcome strata 2 (sometimes FTC-TP BLQ) and selected the marginal outcome sample sizes to be [50, 200, 50]. These target numbers were spread proportionately across dropout strata. To evaluate the performance of BUILD, relative to naïve ODS, 500 random draws from the data were analyzed and compared against the pattern mixture model results from the full data.

### 5.3 Results

Proof of non-ignorable missing effects depends on strong and untestable assumptions. However, assuming the correctness of the chosen model, in the analysis of the full data, a likelihood ratio test for the dropout parameters supported that the dropout mechanism was an informative component of the model (LRT = 29.74, DF = 4, P *<* 0.001) (Fairclough, 2010). Inspection of the estimated parameters suggest that there is a trend toward higher rates of FTC-TP BLQ as study time progresses for those with detectable TFV-DP, which is steeper in those with early dropout. Participants TFV-DP BLQ at baseline begin with a substantially greater risk of FTC-TP BLQ early in time which then begins to improve but stays higher, than those with detectable baseline TFV-DP, for the duration of the study (Figure 2).

**Figure 2.**
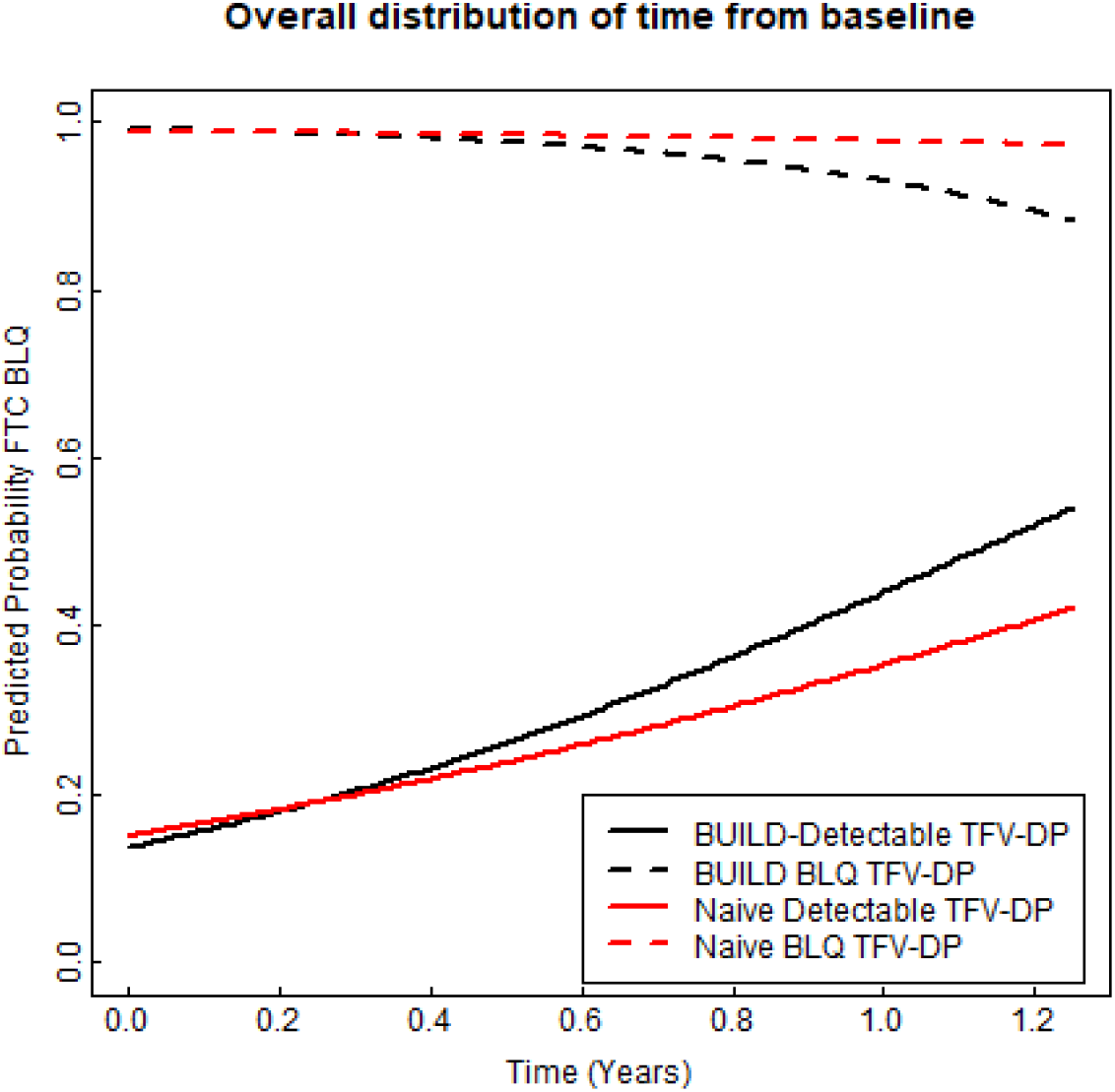
Predicted trajectories in probability of undetectable FTC-TP, by steady state TFV-DP detectable status

After computing marginal parameter estimates, both the time and time by exposure parameters were noticeably different when dropout was taken into account, but comparatively little difference was noted for the model intercept and effect of exposure at baseline (Table 4). While the effect of dropout was not strong enough to change our conclusions about which factors were important contributors to having an undetectable FTC-TP, the magnitudes of the estimates were affected. After considering dropout, the estimated odds of having an undetectable FTC-TP increase by a factor of 5.0 after one year (95% CI: 3.4 - 7.4) among those with detectable TFV-DP at baseline, vs. a reduction in odds by a factor of 0.16 over the same period for those with undetectable TFV-DP at baseline (95% CI: 0.04 - 0.62). In the dropout naïve model the one-year change in estimated odds is 3.09 (95% CI: 2.3 - 4.16) and 0.45 (95% CI: 0.14 - 1.48), respectively.

**Table 4.** Estimates from analyses on the full data, log-odds scale, for both the dropout adjusted and naïve models.

| Parameter | Estimate | Std. Err. | P-value |
| --- | --- | --- | --- |
| Pattern Mixture Parameters |  |  |  |
| Intercept | -1.842 | 0.144 | < 0.001 |
| Time | 1.132 | 0.155 | < 0.001 |
| TFV-DP BLQ | 7.008 | 0.639 | < 0.001 |
| Time * TFV-DP BLQ | -2.245 | 0.737 | 0.002 |
| Early dropout | 0.259 | 0.381 | 0.5 |
| Early dropout * time | 4.161 | 1.218 | < 0.001 |
| Early dropout * TFV-DP BLQ | -1.393 | 0.892 | 0.12 |
| Early dropout * time * TFV-DP BLQ | -7.305 | 2.219 | < 0.001 |
| Dropout Adjusted Parameters |  |  |  |
| Intercept | -1.842 | 0.144 | < 0.001 |
| Time | 1.605 | 0.199 | < 0.001 |
| TFV-DP BLQ | 6.698 | 0.601 | < 0.001 |
| Time * TFV-DP BLQ | -3.868 | 0.762 | < 0.001 |
| Dropout Naïve Parameters |  |  |  |
| Intercept | -1.728 | 0.134 | < 0.001 |
| Time | 1.13 | 0.151 | < 0.001 |
| TFV-DP BLQ | 6.32 | 0.499 | < 0.001 |
| Time * TFV-DP BLQ | -1.929 | 0.633 | 0.002 |

Of the 500 simulation samples 4 BUILD WL, 3 ODS WL models and no ACML models were non-convergent. Model fits were checked for singular variance components, with none found. Consistent with the full data analysis, there was little observed bias in the estimated regression coefficient for the TFV-DP BLQ indicator. BUILD models continued to show little bias in time and time by TFV-DP BLQ interaction estimates. Conversely, ODS only models were substantially biased in time and time by TFV-DP BLQ interaction estimates. ODS only models showed lower coverage and larger MSE in time and time by TFV-DP BLQ interaction estimates while BUILD models held coverage at or above nominal levels. Other metrics were comparable between models (Table 5).

**Table 5.**
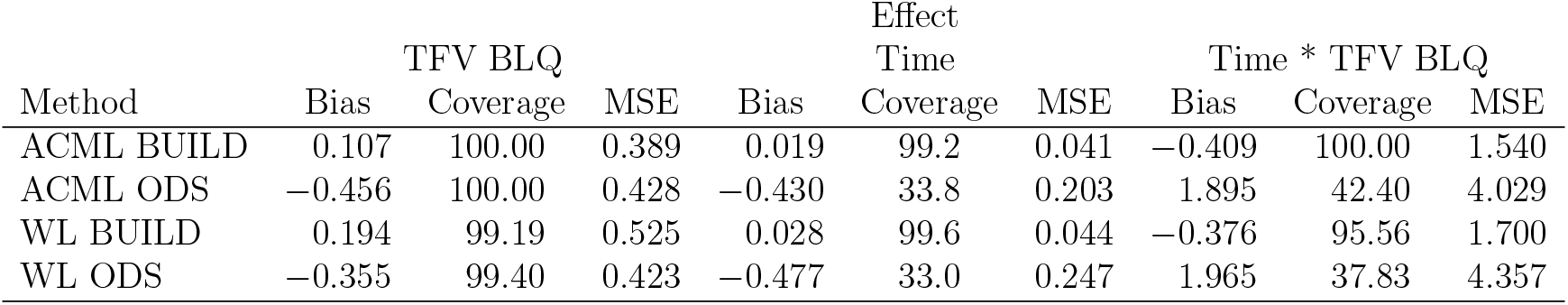
Bias, Coverage and MSE of BUILD and ODS only vs. Full Data Analysis. Results based on 500 random draws of BUILD designs with marginal outcome strata sizes of [50, 200, 50] with proportional distribution of the sample over dropout time.

| Method | TFV BLQ |  |  | Effect<br>Time |  |  | Time * TFV BLQ |  |  |
| --- | --- | --- | --- | --- | --- | --- | --- | --- | --- |
|  | Bias | Coverage | MSE | Bias | Coverage | MSE | Bias | Coverage | MSE |
| ACML BUILD | 0.107 | 100.00 | 0.389 | 0.019 | 99.2 | 0.041 | -0.409 | 100.00 | 1.540 |
| ACML ODS | -0.456 | 100.00 | 0.428 | -0.430 | 33.8 | 0.203 | 1.895 | 42.40 | 4.029 |
| WL BUILD | 0.194 | 99.19 | 0.525 | 0.028 | 99.6 | 0.044 | -0.376 | 95.56 | 1.700 |
| WL ODS | -0.355 | 99.40 | 0.423 | -0.477 | 33.0 | 0.247 | 1.965 | 37.83 | 4.357 |

## 6. Discussion

Longitudinal studies have the potential for nonignorable dropout. Corrected analyses are available and well developed with exposure based sampling and randomized designs, but not for informed sampling strategies. Our proposed expansion to outcome dependent sampling to accommodate dropout effects was tested in both simulated and actual study data and found to minimize bias, hold mean square error to competitive levels, and to maintain coverage probabilities at or above nominal levels. The use of mixture models allows a solution that does not require distributional assumptions on the dropout distribution (Verbeke and Molenberghs, 2000) and is readily extensible to cover a wide variety of dropout associations and even multiple dropout reasons (Moore et al., 2019; Moore et al., 2017, 2019, 2020). In this work we implemented pattern mixture (Little, 1993) and conditional linear models (Wu and Bailey, 1989) to adjust for dropout associations, however, use of penalized splines (Hogan et al., 04a,b) and natural cubic splines (Forster et al., 2011) is also easily implemented. Our BUILD methodology has been encapsulated in an easy to use, and efficient, R package which may be downloaded from the first author’s GitHub page (https://github.com/csevick/build). The question of how to determine optimal sampling probabilities has been left unanswered, but this will be rectified in future work by our team.

The proliferation and increased utilization of stored biosamples is of great importance to biomedical research. The finite nature of these valuable resources demands that we sample as efficiently as possible and analyze results to minimize potential biases whenever possible. BUILD is an adaptable and accessible solution to this problem.

## Supporting information

Supplemental Simulation Results

## Acknowledgements

Dr. Peter L. Anderson received support provided by the grant NIH, NIAID U01 AI084735. This grant funded the iPrEx OLE project, which was the data source for our application example. Thank you to Dr. Diane Fairclough, Dr. Miriam L. Dickinson, and Dr. Elizabeth Juarez-Colunga for helpful comments and review.

A special thank you to Dr. Diane Fairclough, who was always so selfless in helping those around her to grow. Her influence on our research group, both professionally and culturally, cannot be understated. She was a great driver of excellence, creativity, productivity, and the human quality that made our academic community such a supportive and inspiring place. Her passing was a great loss to us all and she is deeply missed.

