## Supplemental Simulation Results for "Statistical Methods for Binary Outcomes Adjusting for Outcome Dependent Sampling in Longitudinal Studies with Nonignorable Dropout"

### 1 Introduction

Due to space constraints, graphical, and partial tabular, results from our simulations (section 4) and application example (section 5) are presented here. Figures are organized by simulation settings and include evaluations of bias, coverage rates and mean square error. A full description of the simulation and analytic methods may be found in the manuscript.

### 1.1 Simulation Results for Dropout Effect on Time only

Table S1: Results based on 500 simulations of BUILD designs with marginal outcome strata sizes of [50, 700, 50] with proportional distribution of the sample over dropout time. Dropout was uniform over study time, beginning at visit 3. 30 imputations used for MI.

| | Target value | | | Bias | | Coverage | | MSE | |
|---|---|---|---|---|---|---|---|---|---|
| | N Follow Up | | | N Follow Up | | N Follow Up | | N Follow Up | |
| Effect | 5 | 10 | Method | 5 | 10 | 5 | 10 | 5 | 10 |
| Intercept | 1.39 | 1.39 | ACML BUILD | 0.009 | 0.007 | 97.2 | 96.2 | 0.017 | 0.015 |
| | | | ACML ODS | 0.005 * | 0.066 | 97.2 | 93.6 | 0.016 | 0.017 |
| | | | MI BUILD | 0.006 | 0.007 | 93.8 | 94 * | 0.014 * | 0.011 * |
| | | | MI ODS | 0.14 | 0.135 | 78.8 | 73 | 0.035 | 0.029 |
| | | | WL BUILD | 0.008 | 0.003 * | 99.2 | 98.4 | 0.017 | 0.015 |
| | | | WL ODS | 0.143 | 0.139 | 94 * | 91.4 | 0.039 | 0.033 |
| Exposure | 0 | 0 | ACML BUILD | 0.012 | 0.008 * | 96.4 | 94 | 0.041 | 0.032 |
| | | | ACML ODS | 0.007 | 0.01 | 96.4 | 94.2 | 0.035 * | 0.025 * |
| | | | MI BUILD | 0.011 | 0.009 | 93.6 | 92.6 | 0.051 | 0.038 |
| | | | MI ODS | 0.006 | 0.015 | 93.8 | 93.2 | 0.055 | 0.036 |
| | | | WL BUILD | 0.012 | 0.009 | 96.4 | 94.2 | 0.052 | 0.038 |
| | | | WL ODS | 0.003 * | 0.013 | 95.6 * | 94.4 * | 0.053 | 0.036 |
| Slope in control | -2.40 | -2.20 | ACML BUILD | 0.021 | 0.004 * | 96 | 95 * | 0.096 | 0.108 |
| | | | ACML ODS | 0.93 | 1.099 | 0.2 | 0 | 0.908 | 1.244 |
| | | | MI BUILD | 0.005 * | 0.021 | 95.4 * | 95.2 | 0.066 * | 0.061 * |
| | | | MI ODS | 1.02 | 1.14 | 0 | 0 | 1.073 | 1.324 |
| | | | WL BUILD | 0.021 | 0.007 | 96.6 | 95 * | 0.106 | 0.107 |
| | | | WL ODS | 1.037 | 1.152 | 0.4 | 0 | 1.128 | 1.367 |
| Slope in exposed | -1.90 | -1.70 | ACML BUILD | 0.001 * | 0.047 * | 95.4 * | 97.8 * | 0.098 * | 0.095 * |
| | | | ACML ODS | 0.988 | 1.131 | 0.4 | 0 | 1.019 | 1.316 |
| | | | MI BUILD | 0.003 | 0.051 | 96.6 | 98 | 0.102 | 0.098 |
| | | | MI ODS | 1.101 | 1.183 | 0 | 0 | 1.241 | 1.425 |
| | | | WL BUILD | 0.003 | 0.051 | 96.6 | 98 | 0.102 | 0.098 |
| | | | WL ODS | 1.1 | 1.184 | 0.4 | 0 | 1.263 | 1.442 |
| time:exposure | 0.50 | 0.50 | ACML BUILD | 0.023 * | 0.056 | 96.2 | 95 * | 0.173 | 0.205 |
| | | | ACML ODS | 0.058 | 0.032 * | 96 | 95.4 | 0.09 * | 0.074 * |
| | | | MI BUILD | 0.024 | 0.065 | 95.8 * | 94.6 | 0.208 | 0.222 |
| | | | MI ODS | 0.081 | 0.043 | 96.2 | 93.2 | 0.109 | 0.085 |
| | | | WL BUILD | 0.024 | 0.064 | 95.8 * | 96 | 0.222 | 0.227 |
| | | | WL ODS | 0.063 | 0.032 | 96 | 94.2 | 0.117 | 0.089 |

*Note:*
* Best performing metric for the group.

### 1.2 Full Simulation Graphical Results

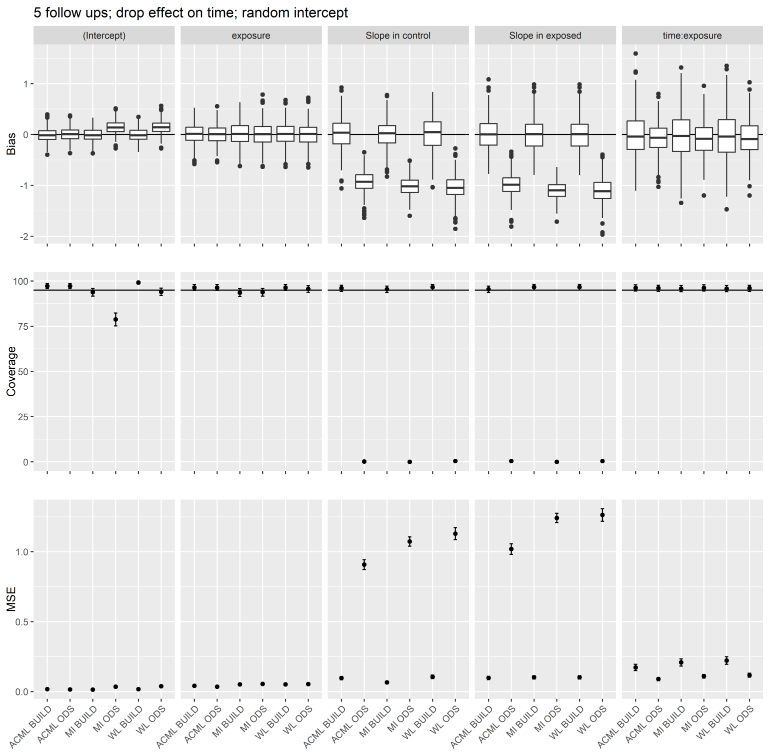


Figure 1: Simulation with 5 follow up points, dropout effect on time, random intercept only. Results based on 500 simulations of BUILD designs with marginal outcome strata sizes of [50, 700, 50] with proportional distribution of the sample over dropout time. Dropout was uniform over study time, beginning at visit 3. 30 imputations used for MI.

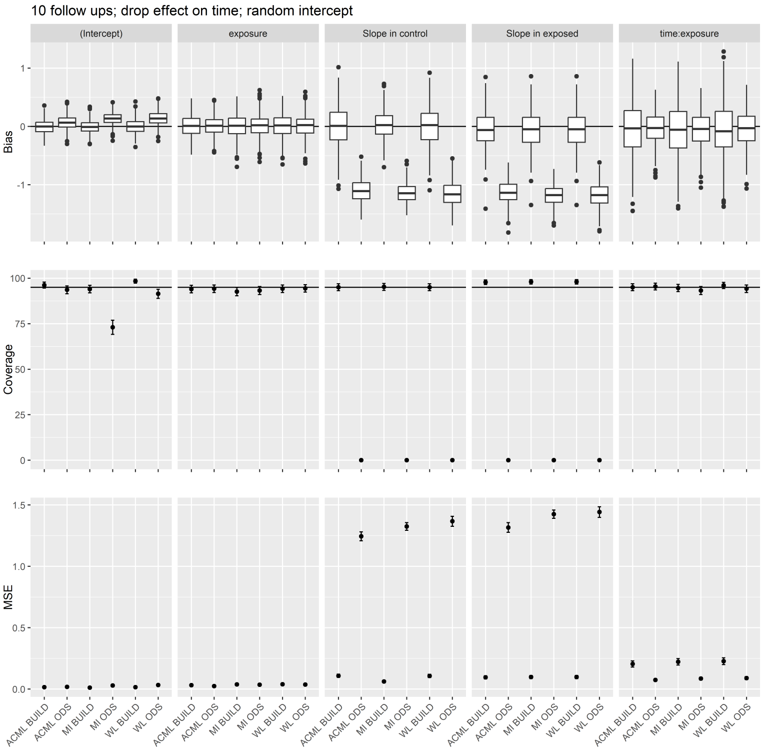


Figure 2: Simulation with 10 follow up points, dropout effect on time, random intercept only. Results based on 500 simulations of BUILD designs with marginal outcome strata sizes of [50, 700, 50] with proportional distribution of the sample over dropout time. Dropout was uniform over study time, beginning at visit 3. 30 imputations used for MI.

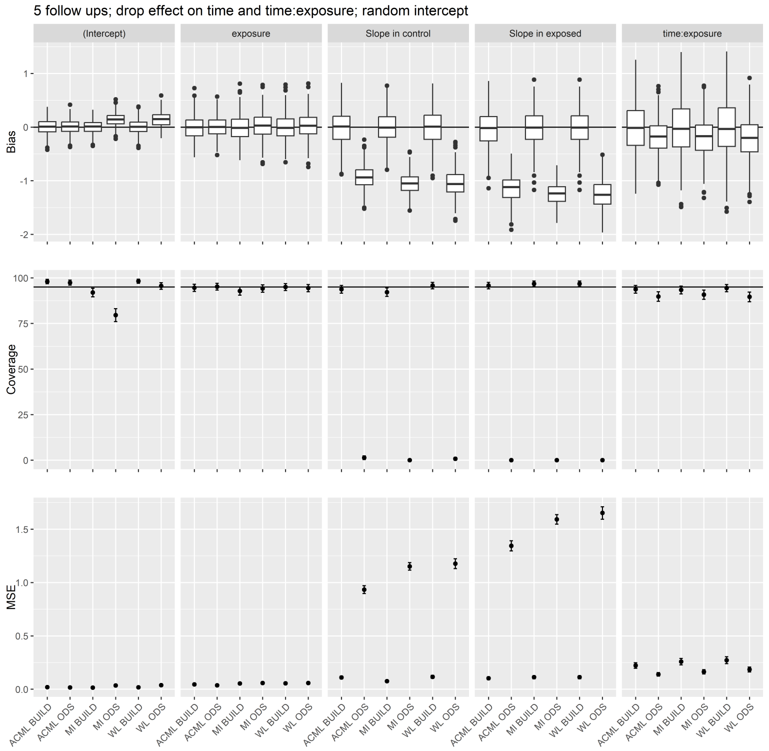


Figure 3: Simulation with 5 follow up points, dropout effect on time and time-exposure, random intercept only. Results based on 500 simulations of BUILD designs with marginal outcome strata sizes of [50, 700, 50] with proportional distribution of the sample over dropout time. Dropout was uniform over study time, beginning at visit 3. 30 imputations used for MI.

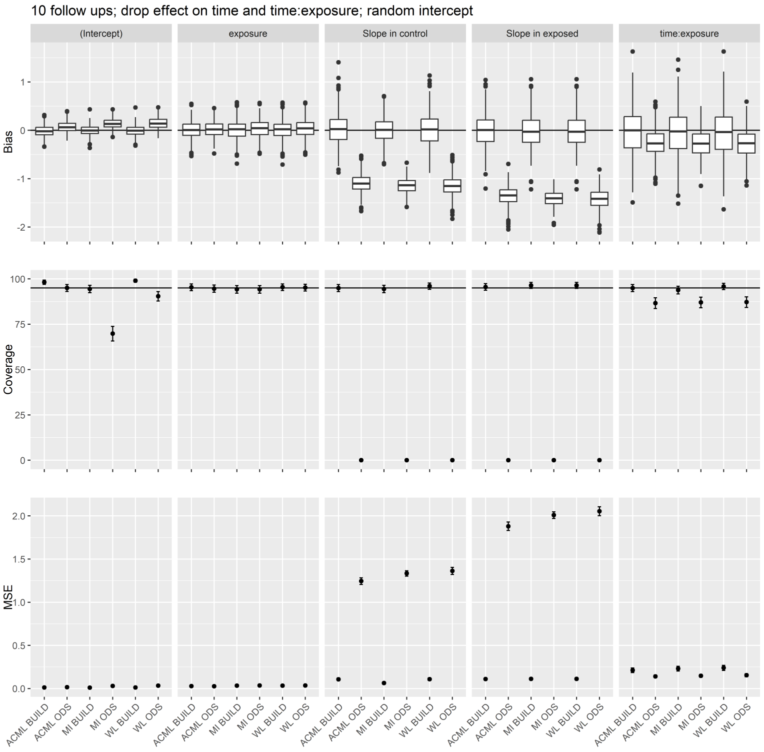


Figure 4: Simulation with 10 follow up points, dropout effect on time and time-exposure, random intercept only. Results based on 500 simulations of BUILD designs with marginal outcome strata sizes of [50, 700, 50] with proportional distribution of the sample over dropout time. Dropout was uniform over study time, beginning at visit 3. 30 imputations used for MI.

### 1.3 Application graphical results

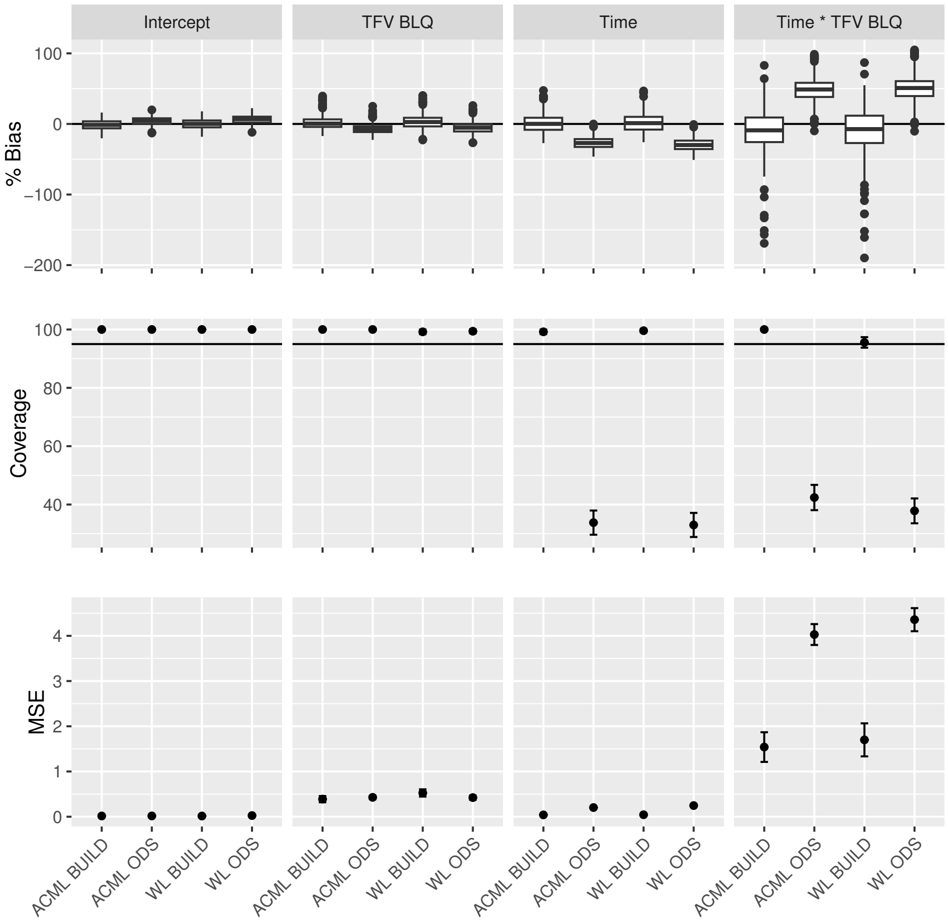


Figure 5: Application Analysis: Bias, Coverage and MSE of BUILD and ODS only vs. Full Data Method
